# JPT2/HN1L functions as an NAADP-binding protein in a cell-type-specific manner

**DOI:** 10.1101/2025.11.11.687795

**Authors:** Roger Ottenheijm, Kai Winterberg, Vivien Throm, Michelle Malz, Merima Bukva, Volodymyr Tsvilovskyy, Rebekka Medert, Andreas H. Guse, Marc Freichel

**Author notes:** Correspondence to: Marc Freichel, INF-366, D-69120 Heidelberg, Germany.

## Abstract

Nicotinic acid adenine dinucleotide phosphate (NAADP) is a second messenger evoking Ca^2+^ release from intracellular Ca^2+^ stores by targeting several Ca^2+^ channels, including two-pore channels (TPC1/2), transient receptor potential mucolipin-1 (TRPML1), or ryanodine receptor type 1 (RYR1). For activation of Ca^2+^ channels, NAADP requires binding proteins, such as JPT2/HN1L and LSM12. So far, their function has been analyzed in several cell lines and only a very limited number of primary cells; however, their physiological relevance in cell types known to utilize NAADP signaling remains unclear. Here, we generated *Jpt2/Hn1l*^*–/–*^ mice to evaluate the contribution of JPT2/HN1L proteins to platelet aggregation and Ca^2+^ signaling in cardiomyocytes, mast cells, and T cells. NAADP is known to contribute to collagen-related peptide (CRP-XL)- evoked platelet aggregation, but this was not altered by JPT2/HN1L deletion. Functional Ca^2+^ imaging revealed that JPT2/HN1L plays a strikingly cell-type specific role in NAADP-mediated Ca^2+^ release. In electrically paced ventricular cardiomyocytes, β-adrenergic stimulation is known to evoke arrhythmogenic spontaneous diastolic Ca^2+^ transients, which were not altered in their frequency in *Jpt2/Hn1l*^*–/–*^ myocytes. Further, antigen-evoked Ca^2+^ transients in peritoneal mast cells (PMCs) are not changed in *Jpt2/Hn1l*^*–/–*^ PMCs. However, CD4^+^ T cells displayed a pronounced requirement for JPT2/HN1L. Following T cell receptor/CD3 stimulation, global Ca^2+^ elevations and early NAADP-driven Ca^2+^ microdomains, which occur within tens of milliseconds of TCR/CD3 engagement and serve as initiating signals for downstream immune activation, were significantly decreased in *Jpt2/Hn1l*^*–/–*^ CD4^+^ cells. We conclude that JPT2/HN1L is indispensable for NAADP-mediated Ca^2+^ release in T cells, but dispensable in cardiomyocytes, platelets, and mast cells, at least for the agonists employed. Accordingly, LSM12 might compensate for the loss of JPT2/HN1L. Together, JPT2/HN1L is not universally required as an NAADP-binding protein but exhibits cell-type specificity, with an essential function in T cell Ca^2+^ signaling.

## Introduction

Nicotinic acid adenine dinucleotide phosphate (NAADP) was first described in 1995 as a highly potent Ca^2+^ releasing second messenger in sea urchin egg homogenate. Afterwards, NAADP was also shown to evoke Ca^2+^ release in several eukaryotic cell types, demonstrating its universal role as a Ca^2+^-mobilizing second messenger. NAADP has been characterized as a potent Ca^2+^- mobilizing agent in mammalian cells, influencing Ca^2+^ release from intracellular stores and playing pivotal roles in cellular signaling across diverse cell types (Churamani et al., 2004; Ruas et al., 2015).

Although NAADP has been identified as a universal Ca^2+^-mobilizing second messenger, its function exhibits specificity within distinct cell types, including T cells (Berg et al., 2000; Dammermann et al., 2009), neurons (Hermann et al., 2020), mast cells (Arlt et al., 2020), cardiomyocytes (Capel & Terrar, 2015), and platelets (Rosado, 2011). In T cells, NAADP signaling contributes to the formation of Ca^2+^ microdomains that are crucial for initiating immune responses (Diercks et al., 2018; Gu et al., 2021; Roggenkamp et al., 2021). In cardiomyocytes, β-adrenergic stimulation leads to Ca^2+^ release from endo-lysosomes originating from activation of two-pore channels (TPC2) and followed by amplification via CaMKII and RYR2 at lysosomal/sarcoplasmic reticulum (Lyso-SR) junctions (Calcraft et al., 2009; Zong et al., 2009). This Ca^2+^ release evokes spontaneous diastolic Ca^2+^ transients (SCTs) in cardiomyocytes, serving as the initial event that triggers arrhythmias under catecholaminergic stress (Capel et al., 2015; Nebel et al., 2013). In platelets, NAADP has been reported to release Ca^2+^ from acidic Ca^2+^ stores (e.g., SERCA3-dependent stores) within seconds upon stimulation, thereby mediating platelet activation and aggregation (Coxon et al., 2012; Feng et al., 2020).

The targets of NAADP-induced Ca^2+^ signaling comprise a range of endo-lysosomal and sarcoplasmic/endoplasmic reticulum (ER) channels, most notably two-pore channels (TPC1 and TPC2), transient receptor potential mucolipin-1 (TRPML1), and ryanodine receptor type 1 (RYR1). TPC channels are structurally unique dimers with six transmembrane domains, are expressed at diverse positions within the endo-lysosomal continuum, and act as the primary NAADP-sensitive channels (Lin-Moshier et al., 2014; Morgan & Galione, 2014). The functional significance of TPCs in NAADP signaling is underscored by their role in modulating physiological processes that vary across cell types. TPC1 is typically associated with early endosomes and plays a role in endosomal recycling, whereas TPC2 predominantly localizes to lysosomes and is integral to lysosomal Ca^2+^ signaling and endocytic trafficking (Castonguay et al., 2017; Morgan et al., 2011). Knockout models for TPC1 and TPC2 have elucidated several roles in cellular processes, including arrhythmogenesis and cardiac remodeling under catecholamine stress (Capel et al., 2015; Nebel et al., 2013), as well as the development of cardiac dysfunction after ischemia/reperfusion (Davidson et al., 2015). In mast cells, TPC1 deficiency or blockade augments systemic anaphylaxis and mast cell activity (Arlt et al., 2020).

Even before the discovery of TPCs as NAADP-sensitive Ca^2+^ channels, RYR1 was proposed to be activated by NAADP (Hohenegger et al., 2002). In fact, RYR1’s open probability was increased by NAADP in lipid planar bilayer experiments using highly purified RYR1 in the presence of recombinant HN1L/JPT2 (Krukenberg et al., 2024). Upon TCR/CD3 stimulation, NAADP is formed in Jurkat T-lymphocytes (Gasser et al., 2006). Further, in RYR1^−/−^ T cells, NAADP-initiated Ca^2+^ microdomains were significantly decreased (Diercks et al., 2018). The involvement of RYR1 in NAADP-mediated Ca^2+^ signaling was also demonstrated by a decrease of NAADP-evoked Ca^2+^ signals upon antagonizing NAADP or upon blocking the RYR1 with antibodies (Dammermann et al., 2009; Gerasimenko et al., 2015). Data on pharmacological inhibition of RYR1, gene silencing, and RYR1 knockout T cells have shown that RYR1 is responsible for the formation of initial, short-lived Ca^2+^ microdomains in T cells. These rapid local Ca^2+^ signals within the first 15 seconds upon T cell activation have been characterized as being NAADP-dependent (Guse & Wolf, 2016; Wolf et al., 2015). Furthermore, TRPML1 is a lysosomal NAADP-sensitive Ca^2+^ channel (Galione, 2011; F. Zhang et al., 2009). The NAADP-evoked Ca^2+^ rise is absent in MEF cells from *Trpml1*^*–/–*^ mice, indicating that TRPML1 channels are a third potential NAADP target (F. Zhang et al., 2011).

A fundamental aspect of NAADP signaling is the requirement for NAADP-binding proteins, as TPC channels are not necessary for NAADP binding (Ruas et al., 2015), and NAADP has never been shown to bind directly to RYR1 or TRPML1. Recently, two key proteins, JPT2/HN1L and LSM12, were discovered as NAADP-binding proteins that facilitate NAADP’s interaction with these channels. JPT2/HN1L was identified via NAADP photoaffinity labeling, protein purification, and mass spectrometry as a potential NAADP-binding protein. JPT2/HN1L deletion in Jurkat T lymphoma cells and primary rat effector T cells results in attenuated Ca^2+^ signals following TCR/CD3 stimulation (Roggenkamp et al., 2021); in parallel, JPT2/HN1L was also identified in red blood cell (RBC) and U2OS cells as an NAADP-binding protein (Gunaratne et al., 2021). LSM12, identified through proteomic approaches as a binding partner of both TPC2 and NAADP, has been shown to modulate NAADP-dependent Ca2+ release with similar capacity, with reduced Ca^2+^ signaling observed in LSM12-deficient cells (J. Zhang et al., 2021). Despite this progress, the roles of these binding proteins in specific cellular contexts of NAADP signaling remain to be elucidated. In this study, we generated JPT2/HN1L-deficient (*Jpt2/Hn1l*^*–/–*^) mice, which enabled us to compare the role of JPT2/HN1L in four primary cell types, for which NAADP signaling has been reported to be triggered after stimulation with (patho)physiological relevant agonists, i.e. platelets, ventricular cardiomyocytes, mast cells and CD4+ T cells from both *Jpt2/Hn1l*^*–/–*^ mice and littermate controls. Our results show that JPT2/HN1L deletion reduces TCR/CD3-mediated global Ca^2+^ elevations and early NAADP-driven Ca^2+^ microdomains. In contrast, NAADP-dependent Ca^2+^ signaling in ventricular myocytes and mast cells, as well as platelet aggregation, were unaltered. These results thus show that the relevance of JPT2/HN1L is cell-type-dependent and suggest a complementary role for other NAADP-binding proteins, possibly LSM12.

## Results

### Generation of *Jpt2/Hn1l*^−/−^ mice

To investigate the role of JPT2/HN1L in primary cell types, we used a genome-editing approach in mouse zygotes, using two guide RNAs flanking exon 3 (Fig. 1A), to generate founder mice as described (Medert et al., 2023). Founder animals harboring a *Jpt2/Hn1l* null (-) allele with deletion of exon 3 (143bp, detected by PCR, Fig. 1B), resulting in a premature stop codon, were bred with C57Bl6/N mice. The obtained heterozygous *Jpt2/Hn1l*^*+/–*^ offspring were intercrossed to *Jpt2/Hn1l*^−/−^ mice (Suppl. Fig. 1A-B). *Jpt2/Hn1l*^*–/–*^ are viable and fertile. Lack of JPT2/HN1L protein expression in the spleen of *Jpt*2/Hn1l–/– mice was confirmed by Western blot analysis (Fig. 1C, Suppl. Fig 1).

**Figure 1.**
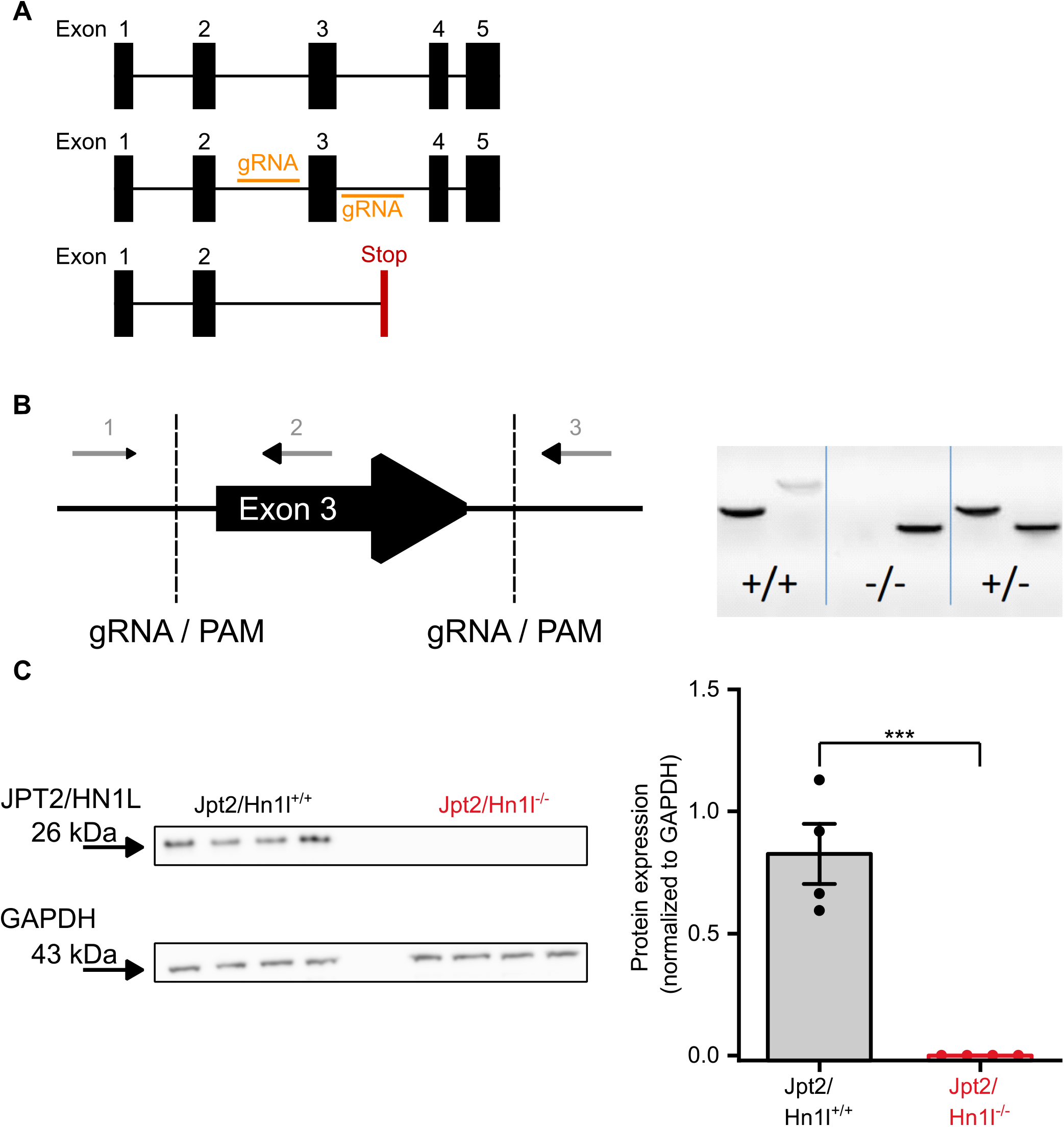
Generation of a Jpt2/Hn1l knockout Mouse. **(A)** cloning strategy of generation of Jpt2/Hn1l knockout mouse using CrisPR-Cas9. Using two gRNAs flanking exon 3 to delete exon 3 and cause a premature stop codon **(B).** PCR to show genetic deletion of exon 3 by Q5-polymerase PCR using primers flanking exon 3. Left reaction (Primer 1 (Jpt2-WT/KO-fw) and 2 (Jpt2-WT-rev2)) results in a WT band of 410bp, and right reaction (primers 1 (Jpt2-WT/KO–fw) and 3 (Jpt2-KO-rev)) shows a knockout band at 299bp and a faint WT band at 900bp. **(C)** Furthermore, deletion of the JPT2 protein in spleens was confirmed by Western Blot using GaPDH as a housekeeping protein. Each datapoint represents protein from a single mouse. Statistical analysis was performed according to the unpaired Student’s t-test.

### JPT2/HN1L does not play a role in catecholamine-induced Ca^2+^ release in cardiomyocytes

Catecholamine stimulation of beta-adrenergic receptors results in NAADP-dependent Ca^2+^ signaling in cardiomyocytes (Lin et al., 2017; Nebel et al., 2013). To test whether JPT2/HN1L plays a role in NAADP-mediated Ca^2+^ signaling in electrically evoked Ca^2+^ transients, we measured the free cytosolic Ca^2+^ concentration ([Ca^2+^]i) in electrically paced cardiomyocytes Next, we Under basal conditions, no difference between SCTs from *Jpt2/Hn1l*^*+/+*^ cells as compared to Jpt2/Hn1l^−/−^ cardiomyocytes was observed (Fig. 2A). Furthermore, the amplitude of Ca^2+^ transients was comparable between these genotypes.

**Figure 2.**
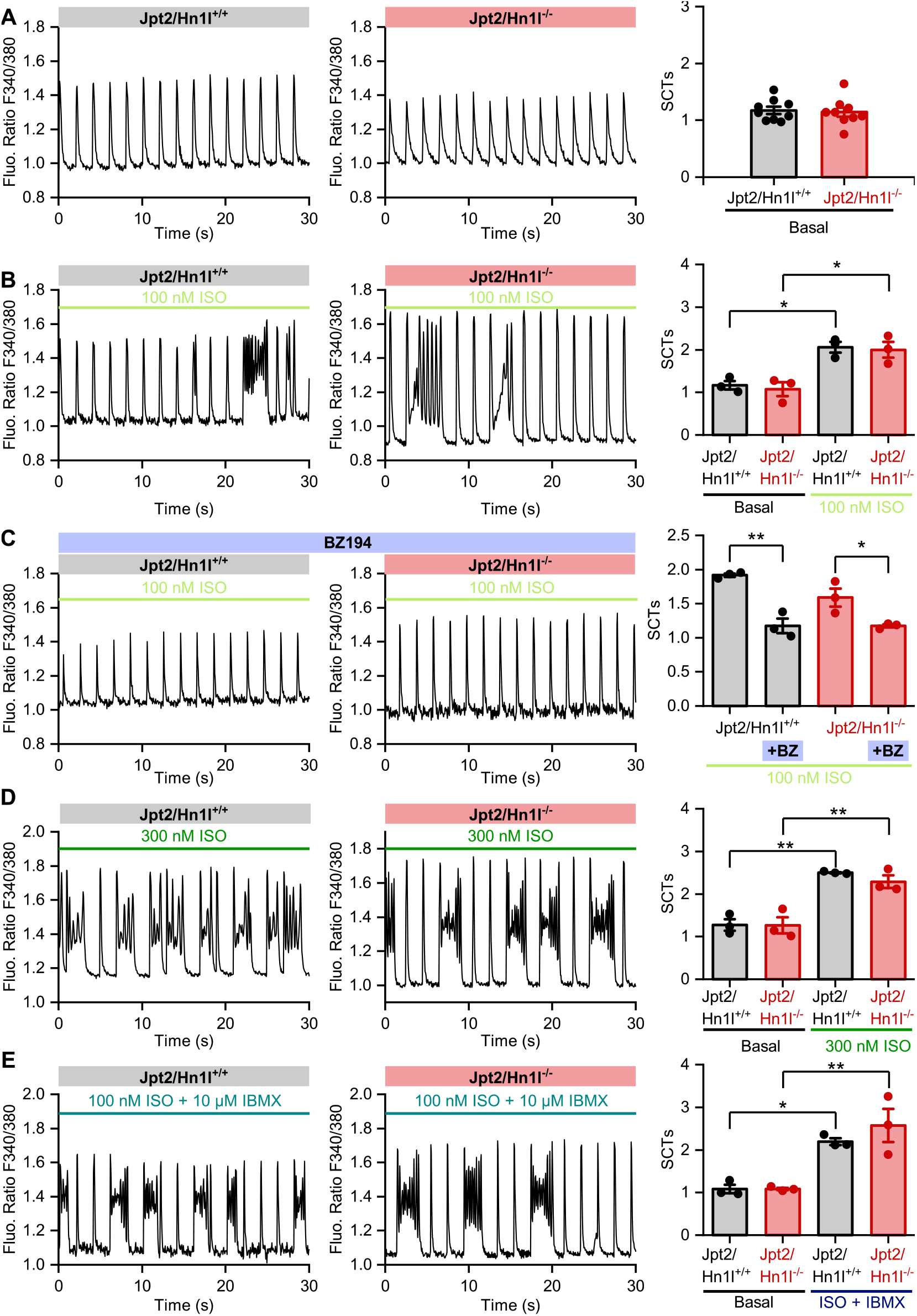
Jpt2/Hn1l does not affect the frequency of spontaneous diastolic Ca^2+^ transients (SCTs) in ventricular cardiomyocytes. **(A)** Pacing (0.5 Hz) ventricular cardiomyocytes with 20Hz acquisition rate evokes Ca^2+^ signals which are independent of Jpt2/Hn1l. **(B)** Application of 100nM ISO evoked increased number of spontaneous Ca^2+^ events in both genotypes. **(C)** ISO-evoked SCTs could significantly blocked by NAADP-antagonist Bz194 (100µM). **(D-E)** Quantification of arrhythmogenic SCTs evoked by **(D)** 300nM ISO or **(E)** combining 100nM isoprenaline with the phosphodiesterase inhibitor IBMX. Each datapoint represents an average of cardiomyocytes of a single mouse and statistical analysis was performed using a Student’s t-test (A-B) or Two-Way ANOVA with Bonferroni Post-hoc testing (C-E).

In cardiomyocytes treated with 100 nM ISO, Ca^2+^ amplitudes increased to 0.46±0.11 (Fura-2 340/380 ratio) and 0.66±0.07 (Fura-2 340/380 ratio) in *Jpt2/Hn1l*^*+/+*^ and *Jpt2/Hn1l*^−/−^ cells, respectively. Furthermore, we observed an increase in the number of SCTs in both *Jpt2/Hn1l*^*+/+*^ and *Jpt2/Hn1l*^−/−^ cells (Fig. 2B, Suppl. Fig. 2). Isoprenaline (ISO) stimulation during electrical pacing evokes NAADP-mediated spontaneous diastolic Ca^2+^ transients (SCT). Pre-incubation of cardiomyocytes with the NAADP-antagonist BZ194 (100 µM) resulted in a significantly reduced number of ISO-evoked SCTs in both *Jpt2/Hn1l*^*+/+*^ and *Jpt2/Hn1l*^−/−^ cells (Fig. 2C), confirming previous results (Nebel et al., 2013).

Further increasing the concentration of ISO to 300nM resulted also in similar number of SCT (Fig. 2D) in Jpt2/Hn1l^+/+^ compared to *Jpt2/Hn1l*^−/−^ and an amplitude in intracellular Ca^2+^ of 0.35±0.04 (Fura-2 340/380 ratio) and 0.39±0.13 (Fura-2 340/380 ratio) in *Jpt2/Hn1l*^*+/+*^ and *Jpt2/Hn1l*^−/−^ cardiomyocytes, respectively (Fig. 2D, Suppl. Fig. 2). 3-Isobutyl-1-methylxanthin (IBMX) is a cAMP-phosphodiesterase (PDE) inhibitor. PDEs catalyze the degradation of cAMP and can be inhibited to prolong its action. cAMP has been suggested to induce Ca^2+^ release which is reduced in *TPC2*^−/−^ cells (Chang et al., 2020). We aimed to further enhance the ISO-evoked Ca^2+^ response by combining ISO with IBMX. Nevertheless, no change in frequency or amplitude of SCT was observed between genotypes (Fig. 2E, Suppl. Fig. 2).

### NAADP-mediated platelet aggregation is unaffected in *Jpt2/Hn1l*^*–/–*^ platelets

Upon activation, platelets undergo morphological changes and form aggregates. This activation is tightly regulated by signaling molecules, particularly cytosolic Ca^2+^ levels (Rosado, 2011). Stimulation of Collagen-Related Peptide (CRP-XL) activates the GP-VI receptor, which has also been postulated to evoke NAADP-dependent Ca^2+^ release and aggregation (Coxon et al., 2012). We stimulated platelets with CRP-XL at a concentration that induced submaximal aggregation and is NAADP-dependent. Platelet aggregation upon stimulation of 0.1µg/ml CRP-XL was unchanged in JPT2/HN1L-deficient platelets compared to litter controls (Fig. 3A). Taken together, these data show that loss of JPT2/HN1L does not alter NAADP-mediated aggregation in mouse platelets. However, 50 µM of the NAADP antagonist Ned19 (Naylor et al., 2009) has previously been shown to reduce CRP-XL-evoked aggregation in human platelets (Coxon et al., 2012). Here, we could show that Ned19 also abolished the 0.1 µg/ml CRP-XL-evoked platelet aggregation in mouse platelets (Fig. 3A).

**Figure 3.**
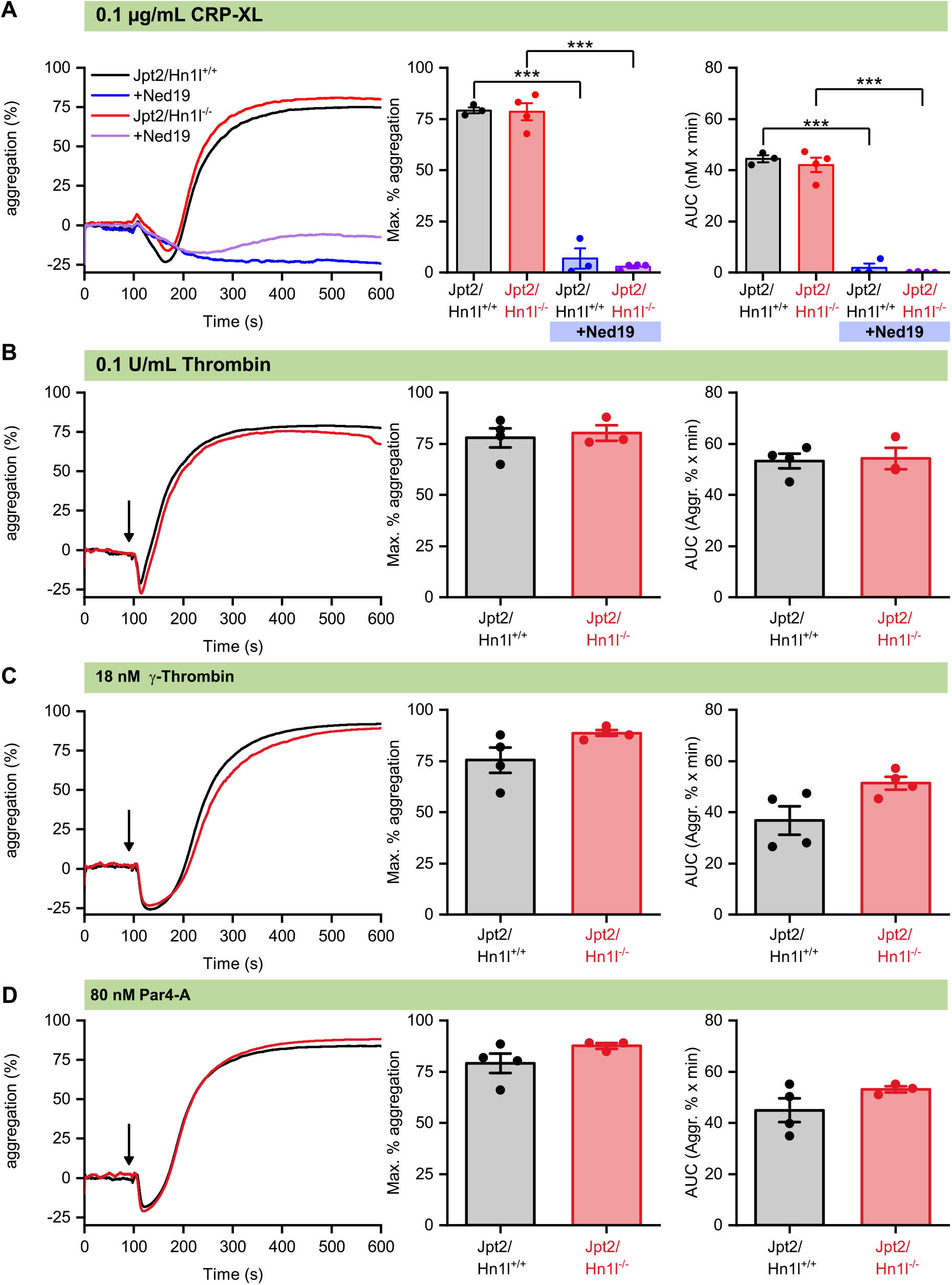
latelet aggregation is independent of Jpt2/Hn1l. **(A)** Platelet aggregation and quantification of maximal aggregation and AUC evoked by Collagen-related peptide (CRP-XL) is unchanged in Jpt2/Hn1l^-/-^ compared to Jpt2/Hn1l^+/+^, and can be significantly inhibited by the NAADP-antagonist Ned19. **(B-D)** Comparison of aggregation evoked by **(B)** thrombin, **(C)** y-thrombin, or **(D)** Par4-A in Jpt2/Hn1l^+/+^ and Jpt2/Hn1l^−/−^ platelets shows no differences in maximal aggregation and AUC. Each data point represents a biological replicate of platelets isolated from a single group of mice. Statistical significance was tested using an unpaired Student’s t-test or two-Way ANOVA.

In addition to GPVI receptors, PAR4 activation has also been described to generate NAADP in platelets. PAR-4 receptors can be stimulated by low concentrations of thrombin (0.1 U/ml) as well as by PAR-4-specific agonists, e.g., y-thrombin or PAR-4A peptides (Jardin et al., 2007; Soslau et al., 2004). For these agonists, we generated dose-response curves to determine IC50 Values for aggregation (Suppl. Fig. 3).

However, aggregation upon stimulation of the PAR-4 receptor using 0.1 U/ml thrombin resulted in a maximal aggregation of and AUC Jpt2/Hn1l^+/+^, which was comparable to that of Jpt2/Hn1l^−/−^ platelets (Fig. 3 B), respectively. We also observed no significant difference in aggregation upon stimulation with the PAR4-specific agonist γ-thrombin in a maximal aggregation and AUC between Jpt2/Hn1l^+/+^ and Jpt2/Hn1l^−/−^ platelets (Fig. 3C). Similarly, PAR4-A, another agonist specifically for the PAR4 receptor, evoked an aggregation in Jpt2/Hn1l^+/+^ platelets which was indistinguishable from Jpt2/Hn1l^−/−^ platelets (Fig. 3D).

### Peritoneal mast cells

To test the impact of the deletion of the NAADP-binding protein JPT2/HN1L on the antigen-evoked Ca^2+^ signaling in PMCs, we sensitized PMCs with anti-IgE antibodies directed against DNP. We studied the DNP-evoked changes in [Ca^2+^]_i_ in PMCs from *Jpt2/Hn1l*^−/−^ and litter-matched *Jpt2/Hn1l*^+/+^ controls. The Ca^2+^ rise evoked by antigen stimulation (DNP 10 ng/ml) did not differ in amplitude or AUC between genotypes (Fig. 4A). However, we could show that DNP-evoked Ca^2+^ rise depends on NAADP, as we were able to significantly reduce DNP-evoked Ca^2+^ rise by pre-incubation with BZ194 (100µM) in both JPT2/HN1L^+/+^ and JPT2/HN1L^-/-^ cells (Fig. 4B).

**Figure 4.**
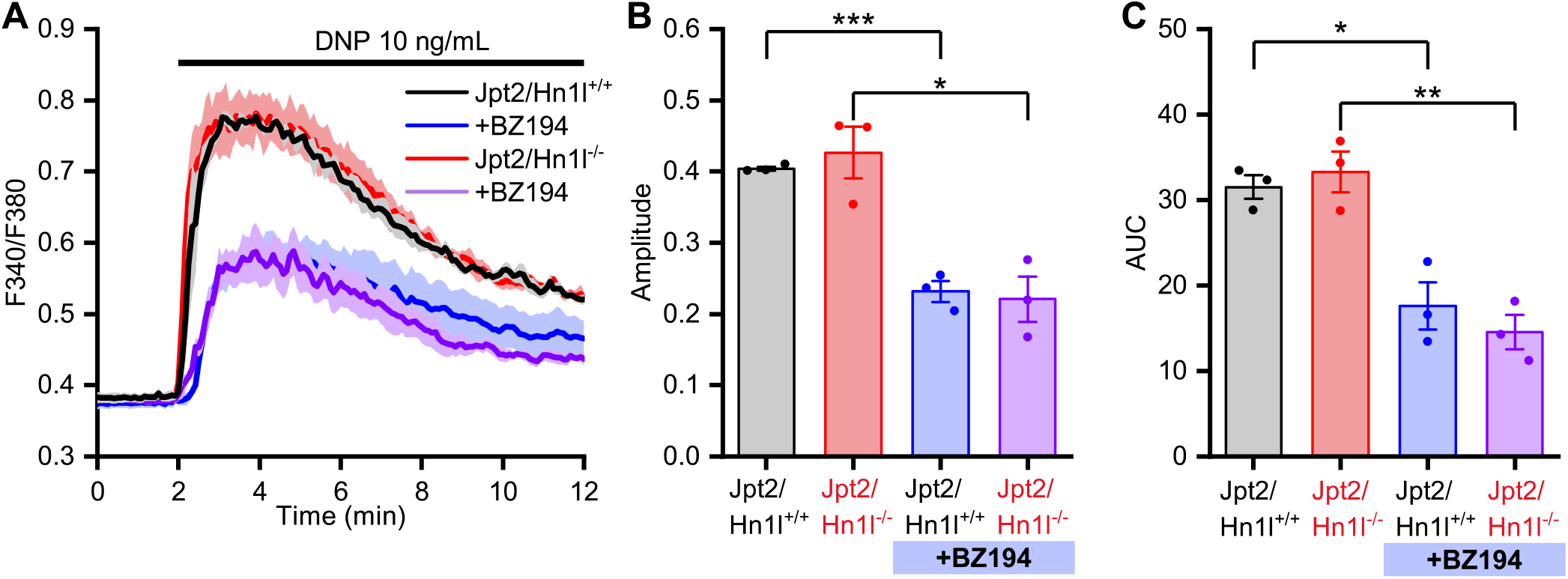
No difference in antigen-induced Ca^2+^ rise in Jpt2/Hn1l-deficient and wild-type PMCs. **(A)** Mean traces of F340/F380 loaded PMCs isolated from Jpt2/Hn1l^−/−^ and litter-matched Jpt2/Hn1l^+/+^ controls representing changes of [Ca^2+^]_i_ stimulated with 10 ng/ml of antigen (DNP) in the presence or absence of the NAADP-antagonist Bz194. **(B, C)** The corresponding statistical analysis of the amplitude (B) and AUC (C). **(D)** Mean traces of changes of [Ca^2+^]_i_ are shown as F340/F380. Statistical significance was tested using an unpaired Student’s t-test.

### JPT2/HN1L is essential for NAADP-dependent Ca^2+^ microdomains and global Ca^2+^ signaling in CD4^+^ T cells

One of the essential signaling events during T cell activation is Ca^2+^ signaling. NAADP serves as a pivotal second messenger, initiating Ca^2+^ signaling events that begin within seconds of TCR/CD3 engagement. Basal Ca^2+^ levels in *Jpt2/Hn1l*^*–/–*^ T cells were unchanged compared to *Jpt2/Hn1l*^*+/+*^ controls (Fig. 5A, B). However, upon TCR/CD3 stimulation, a significant reduction in Ca^2+^ peak was observed in JPT2/HN1L-deficient T cells compared to control cells (Fig. 5A, B). Then, we analyzed the earliest detectable Ca^2+^ signals in T cells, the Ca^2+^ microdomains that occur predominantly beneath the plasma membrane near the immune synapse (Fig. 5C). These Ca^2+^ microdomains form within tens to hundreds of milliseconds of TCR/CD3 ligation and precede global Ca^2+^ increases. In *Jpt2/Hn1l*^−/−^ T cells, a significant decrease in the number of Ca^2+^microdomains was observed, shown here as integrated over the first 15s (Fig. 5C), or as kinetics data (Fig. 5D). Together, these data confirm the relevance of JPT2/HN1L in NAADP-mediated Ca^2+^ release in CD4^+^ T cells.

**Figure 5.**
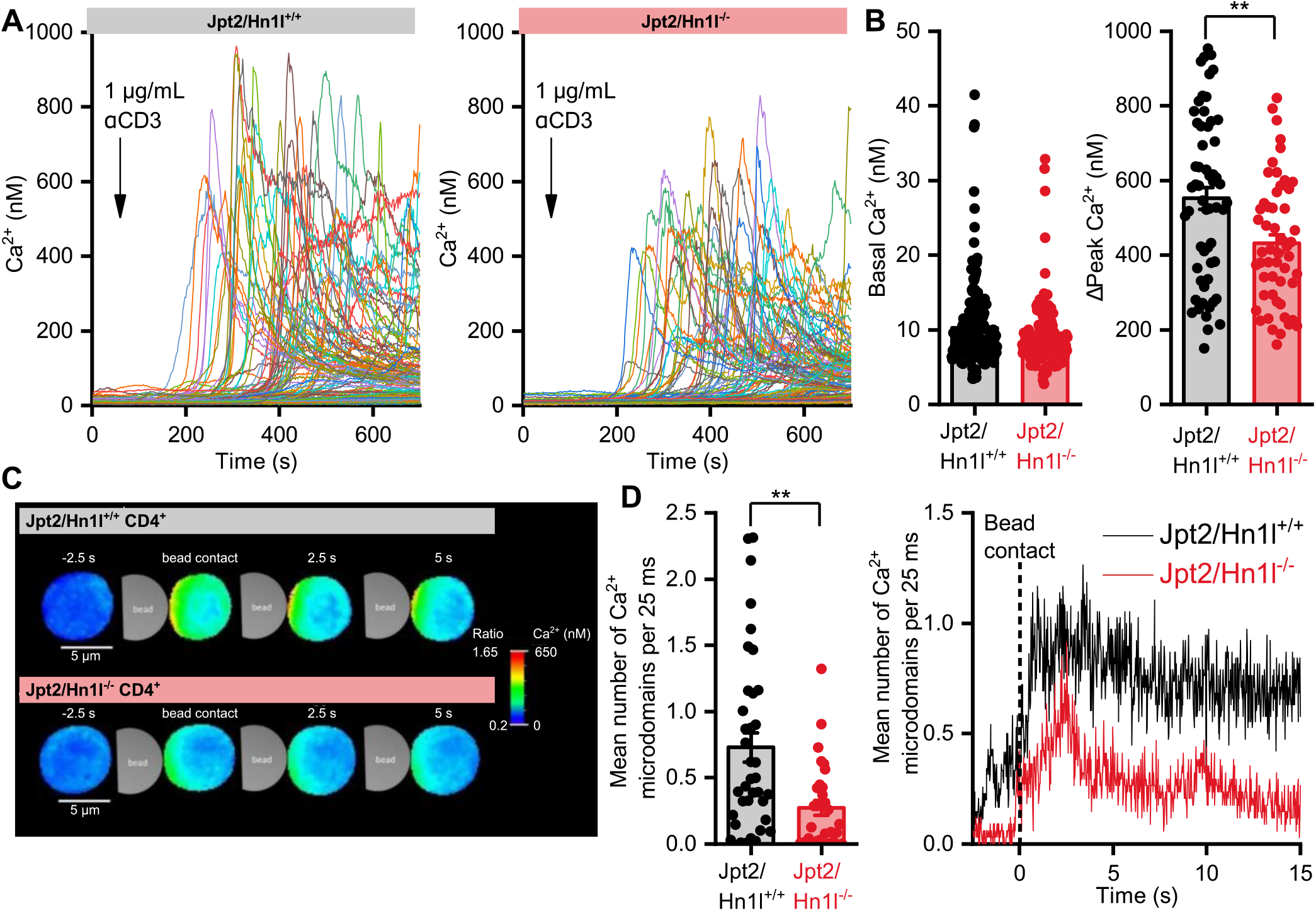
Global and local Ca^2+^ signals in CD4+ T cells. **(A, B)** Analysis of global Ca^2+^ signals in CD4^+^ T cells loaded with Fura2. **(A)** Whole cell Ca^2+^ traces after addition of 1 µg/mL soluble anti-CD3ε antibody at 60s. Images were acquired at 0.5 fps, n = 179 *Jpt2/Hn1l*^*+/+*^ cells (left panel) and 150 *Jpt2/Hn1l*^*–/–*^ cells (right panel) from N = 3 spleens per genotype. **(B)** Data points and average basal Ca^2+^ level and peak Ca^2+^ increase measured in Fura2-loaded T cells after anti-CD3ε treatment. 58 peaks detected in *Jpt2/Hn1l*^*+/+*^ cells and 53 peaks detected in *Jpt2/Hn1l*^*–/–*^ cells were compared with an unpaired t-test. **(C, D)** Analysis of Ca^2+^ microdomains in CD4^+^ T cells. **(C)** Maximum intensity projections of 5 frames from representative cells showing Ca^2+^ microdomains at the bead contact site. The cells were loaded with Fluo4 and FuraRed and stimulated with anti-CD3ε/anti-CD28 beads for subcellular ratiometric Ca^2+^ imaging. Images were acquired at 40 fps. **(D)** Number of Ca^2+^ microdomains in 15s recording period (left) and average time course of Ca^2+^ microdomains before and after bead contact (right) from n = 37 *Jpt2/Hn1l*^*+/+*^ and 31 *Jpt2/Hn1l*^*–/–*^ cells derived from N = 6 spleens per genotype, statistical analysis with Mann-Whitney U test. **: p < 0.01, ***: p < 0.001.

## Discussion

In this study, we generated a JPT2/HN1L knockout mouse and systematically assessed the involvement of JPT2/HN1L as an NAADP-binding protein in four primary cell types (cardiomyocytes, platelets, mast cells, and CD4^+^ T cells). All these cell types have previously been described as utilizing NAADP as a second messenger for Ca^2+^ mobilization from intracellular stores. Our findings have provided new insights into the physiological relevance of JPT2/HN1L, demonstrating apparent cell-type specificity in its role in NAADP-mediated Ca^2+^ signaling. JPT2/HN1L is not required for catecholamine-induced Ca^2+^ release in ventricular cardiomyocytes and for NAADP-dependent platelet aggregation. Similarly, NAADP-mediated Ca^2+^ release in mast cells upon DNP stimulation was unaffected by JPT2/HN1L deletion. However, it was critical for the generation of early NAADP-driven Ca^2+^ microdomains in murine CD4^+^ T cells.

Previous studies have implicated NAADP signaling in myocardial Ca^2+^ regulation, particularly during β-adrenergic stress and arrhythmogenesis (Capel et al., 2015; Nebel et al., 2013). NAADP has been shown to stimulate Ca^2+^ release mediated by TPC2 channels at lysosomal-sarcoplasmic reticular junctions of cardiomyocytes. Our data, however, demonstrate that loss of JPT2/HN1L did not affect Ca^2+^ transients under basal pacing or following ISO stimulation, even when PDE inhibition potentiated the response. The frequency and amplitude of spontaneous Ca^2+^ transients and SCTs remained similar between *Jpt2/Hn1l*^*+/+*^ and *Jpt2/Hn1l*^*–/–*^ cardiomyocytes across multiple conditions. Platelets are activated upon increased Ca^2+^ signaling, and NAADP has been proposed to contribute to platelet activation downstream of PAR-4 and GPVI receptors (Coxon et al., 2012; Rosado, 2011). Importantly, pharmacological inhibition of NAADP with Ned19 partially impairs collagen-related peptide–induced aggregation, supporting NAADP’s role as an intracellular amplifier. Our results, however, show that platelet aggregation triggered by low-dose thrombin, γ-thrombin, PAR-4 peptide, or CRP-XL was preserved in JPT2/HN1L-deficient platelets, with no significant differences compared to controls. Finally, DNP has been shown to induce NAADP-mediated Ca^2+^ release via TPCs in mast cells; however, these Ca^2+^ signals were unchanged by genetic deletion of JPT2/HN1L. These findings imply that JPT2/HN1L is not required for NAADP-mediated signaling in cardiomyocytes, platelets, and mast cells, at least after stimulation with the agonists used in this study.

This could be explained by cell-type–specific reliance on distinct NAADP-binding proteins, or by the deletion of JPT2/HN1L, which evokes redundant mechanisms. Previous studies have described another NAADP-binding protein, LSM12 (J. Zhang et al., 2021). LSM12 was shown in HEK cells to interact with NAADP, TPC1, and TPC2, and to be involved in NAADP-evoked TPC activation and Ca^2+^ mobilization from acidic Ca^2+^ stores (J. Zhang et al., 2021). Furthermore, both JPT2/HN1L and LSM12 bind NAADP and function as NAADP-binding proteins in U2OS cells. In these cells, genetic deletion of Lsm12 was insufficient to completely suppress NAADP-mediated Ca^2+^ release. It required an additional knockout of JPT2/HN1L (Gunaratne et al., 2023), suggesting the possibility of redundant mechanisms between JPT2/HN1L and LSM12.

In contrast, JPT2/HN1L played a critical role in NAADP-mediated Ca^2+^ signaling in CD4^+^ T cells. Our results are highly consistent with previously published data obtained in the same cell type and host species. Specifically, comparable effect sizes have been reported for the DUOX1/2 double knockout (Gu et al., 2021) and for CD4^+^ T cells lacking RYR1 (Diercks et al., 2018; Tabrizi et al., 2026). Of note, in contrast to cardiomyocytes and mast cells, NAADP triggers Ca^2+^ release in T cells via RYR1. Following TCR/CD3 stimulation, global Ca^2+^ elevations were significantly decreased in *Jpt2/Hn1l*^−/−^ T cells. More markedly, the formation of Ca^2+^ microdomains immediately after TCR/CD3 ligation is a hallmark of NAADP-dependent signaling and was strongly reduced in JPT2/HN1L-deficient T cells. These highly localized Ca^2+^ transients serve as critical initiating events for T cell activation (Diercks et al., 2018; Gu et al., 2021; Roggenkamp et al., 2021; Wolf et al., 2015), leading to downstream global Ca^2+^ signaling, NFAT activation, and gene transcription. Also, in Neuro2A cells, JPT2/HN1L appears indispensable for NAADP-evoked Ca^2+^ release, as Jpt2/Hn1l–/–Neuro2A cells did not respond to stimulation with the membrane-permeant NAADP precursor MASTER-NAADP (Krukenberg et al., 2024).

Our data confirm previous evidence (Diercks et al., 2018; Roggenkamp et al., 2021; Wolf et al., 2015) that JPT2/HN1L is critical for linking NAADP to RYR1-mediated Ca^2+^ release in T cells. Of note, this interaction of RYR1, JPT2/HN1L, and NAADP was directly shown by increased RYR1 open probability when highly purified RYR1 was fused into lipid bilayers, and upon addition of de-esterified MASTER-NAADP and recombinantly expressed and purified JPT2/HN1L (Krukenberg et al., 2024). This suggests that the T cell signaling architecture differs from that of other cell types studied, indicating that NAADP-binding proteins have cell-type-specific functions. One possibility for explaining this difference between cell types might be (i) that the ion channel target of NAADP in CD4^+^ T cells has been identified as RYR1, whereas (ii) in the other cell types tested (cardiomyocytes, platelets, and PMCs), the ion channels involved might rather be TPCs. However, based on our data, we cannot conclude whether this is TPC1 or TPC2, or both. The essential role of JPT2/HN1L in T cell Ca^2+^ signaling also positions it as a potential therapeutic target. Given the importance of early Ca^2+^ events in TCR signaling, modulation of JPT2/HN1L may represent a means of fine-tuning immune activation in settings such as autoimmunity, transplantation, or immunotherapy (Chen et al., 2024; Heßling et al., 2023).

Together, these findings demonstrate that JPT2/HN1L is not universally required as an NAADP-binding protein but instead has cell-type–specific functions. In cardiomyocytes and platelets, JPT2/HN1L deficiency is well tolerated, suggesting that alternative NAADP-binding proteins (such as LSM12) or parallel Ca^2+^ release pathways can compensate. Conversely, T cells are critically dependent on JPT2/HN1L for the generation of early NAADP-driven Ca^2+^ microdomains, highlighting a certain degree of specialization. Further studies should aim to dissect the interplay between JPT2/HN1L and LSM12, to uncover whether compensatory upregulation occurs in specific tissues, and to define the broader network of NAADP-interacting proteins. In initial qPCR experiments, we observed significant compensation in Lsm12 expression in *Jpt2/Hn1l*^−/−^ cardiomyocytes, whereas this was not observed in spleens from *Jpt2/Hn1l*^−/−^ mice, from which we isolated the T cells analyzed in Ca^2+^ imaging experiments (Suppl. Fig. 4). Importantly, our JPT2/HN1L knockout model provides a valuable tool to address these questions in vivo. Ultimately, delineating the mechanisms that confer cell-type specificity to NAADP signaling will be critical to harnessing this pathway for therapeutic interventions in cardiovascular disease, hemostasis, and immune regulation.

## Materials and Methods

### Animal Experiments

Mice were maintained under specified pathogen-free conditions in the animal facility (IBF) of the Heidelberg Medical Faculty. Mice were housed in a 12 h light–dark cycle, with a relative humidity of 56–60%, an air change rate of 15 per hour, and a room temperature of 22±2 °C. They were kept in conventional cages, type II or type II long, provided with animal bedding LTE E-001 (ABBEDD, Germany) and tissue papers as enrichment. Standard autoclaved food (Rod 16 or 18, Altromin, Lage, Germany) and autoclaved water were available to consume ad libitum.

### Generation of the JPT2/HN1L knockout mouse

To generate a JPT2 knockout mouse, we used a genome-editing approach in mouse zygotes, which were injected with heiCas9 mRNA (Thumberger et al., 2022) as previously described (Medert et al., 2023). We designed sgRNAs that induced double-strand breaks at the introns flanking exon 3, causing deletion of the sequences between the sgRNAs, as shown previously (Medert et al., 2023), but without providing a donor sequence, thereby allowing NHEJ-mediated repair. Deletion of exon 3 introduces a premature stop codon in the *Jpt2* coding sequence. Post-injection, embryos were transferred to pseudo-pregnant females, and founders were screened by PCR genotyping to confirm the deletion. Six out of seven founder mice carried the null allele. Founder #6 was crossed to C57Bl6/N mice. Sequencing of DNA fragments amplified from genomic DNA of the *Jpt*2^+/–^ offspring validated the absence of exon 3. Subsequently, all mice were genotyped by collecting an ear punch biopsy at 3 weeks of age. Genotypes were confirmed by PCR. *Jpt*2^+/–^ mice were intercrossed to obtain *Jpt*2^−/−^ offspring.

### Western Blot

Mice were sacrificed, spleens were harvested and homogenised in lysis buffer containing protease inhibitors (1 mM Iodoacetamide; 1 mM Phenanthroline; 0.1 mM PMSF; 1 µg/ml Antipain; 1 µg/ml Leupeptin; 0.7 µg/ml Pepstatin; 1 mM Benzamidine; 0.3 µM Aprotinin) using a tissue grinder (Tenbroeck Tissue Grinders, Wheaton). The BCA assay determined protein concentration. Immunoblotting of microsomal membrane fractions was performed as previously described (Medert et al., 2020). Samples were denatured at 60°C for 20 min and loaded on Bis-Tris Plus gels, followed by gel electrophoresis at 150 V for 60 min. Resolved proteins were blotted onto a 0.45 µm Protran nitrocellulose membrane (GE Healthcare) at 10 V for 60 min. Next, the membrane was blocked in 5% non-fat milk with TBS (pH 7.5). The Blot was incubated with a JPT2 (Sigma, HPA041888) or GAPDH (Acris, ACR001PS) antibody at 4 °C overnight. Subsequently, the blot was washed three times for 5 min in TBST and incubated in horseradish peroxidase-conjugated anti-rabbit (GE Healthcare) and anti-mouse (Sigma) antibodies at 1:50.000, respectively, at room temperature (RT) for 1 h. Afterwards, the blot was washed twice for 5 min in TBST and once for 5 min in TBS and then incubated with SignalFire Elite ECL (Cell Signalling) for 1 min. Chemiluminescence was detected using digital imaging (GE Healthcare, ImageQuant LAS 4000 mini). Original scans were saved as TIFF files from a LAS 4000 mini (GE Healthcare) and further analyzed using FIJI.

### Isolation of adult mouse ventricular cardiomyocytes and intracellular Ca^2+^ measurements

For isolation of adult mouse ventricular cardiomyocytes, retrograde perfusion of coronary arteries was performed as described before (Camacho Londoño et al., 2015). Briefly, the heart of an anaesthetized mouse was removed and cannulated from the aorta with a blunted 27G cannula and fixed by a ligature. First, the heart was washed with an oxygenated EGTA-containing (200 µM) Solution A+ for 3 min at RT. Subsequently, the heart was perfused with oxygenated digesting buffer (Solution A with 0.05 mg/ml Liberase TM, Roche) for 2-3 min at 37°C until it became soft and pale. After digestion, the atria were separated from the ventricle. Ventricular cells were dissociated by gently pipetting, and undissociated tissue was removed by filtration through a 150 µm filter. The extracellular Ca^2+^ concentration was then stepwise increased to 1 mM. Cardiomyocytes were seeded on ECM-coated coverslips in Ca^2+^-containing (1.8 mM) Tyrode’s solution.

To investigate electrically evoked Ca^2+^ transients, Fura2-AM-loaded cardiomyocytes were stimulated using an electrical field stimulator (MyoPacer Cell Stimulator, IonOptix) set to 5 V and 0.5 Hz and custom-made electrodes (Fachabteilung 3.1 Elektronik, Zentralbereich Neuenheimer Feld, Heidelberg University). Cells were continuously perfused with a Ca^2+^-containing buffer throughout the measurement. During the measurements cells were stimulated with 5V at 0.5 Hz. for 5 min, cells were perfused with or without 100 nM ISO in 1.8 mM extracellular Ca^2+^. Spontaneous calcium transients were defined as the number of calcium transients divided by the number of pacing events.

### Platelet isolation and Aggregation

Blood sampling and animal handling were approved and performed in accordance with the regulations of the Regierungspräsidium Karlsruhe, the University of Heidelberg and in conformity with the guidelines of Directive 2010/63/EU of the European Parliament on the protection of animals used for scientific purposes. Blood was obtained by submandibular bleeding as described (Golde et al., 2005) but using an 18G 11/2” needle under terminal Ketamine/Xylazine anesthesia (Xylazine-hydrochloride and Ketamine-hydrochloride final dose 16 and 120 mg/kg body weight, respectively; dissolved in sterile NaCl 0.9% and volume of 5 µL/g body weight; intraperitoneal injection). Washed platelets were isolated similarly as previously reported (Yang et al., 2022). Platelet numbers were determined and adjusted to 3 × 10^8^ platelets/mL, and cells were incubated for 30 min at 37 °C in the presence of 0.02 U/mL apyrase before aggregation experiments. Aggregation was determined at 37 °C using two APACT 4S Plus aggregometers (Diasys, Holzheim, Germany). Tyrode’s buffer served as the reference for 100% light transmission. A 190 µL platelet suspension was used and stimulated with corresponding agonists in a volume of 10 µL after 90 s of basal recording. In the case of ADP-induced aggregation, measurements were performed in the presence of 70 µg/mL fibrinogen (F3879, Sigma-Aldrich, Taufkirchen, Germany), added (4 µL) shortly before starting measurements.

### Peritoneal Mast Cell Isolation and intracellular Ca^2+^ measurements

Peritoneal mast cells (PMCs) were isolated and cultured as previously described (PMID: 30035759). Briefly, the cells were isolated by peritoneal lavage with RPMI medium, centrifuged (300 g), and subsequently re-suspended in 4 ml medium per mouse. The RPMI-based culture medium was supplemented with 20% fetal calf serum (FCS), 1% Penicillin-Streptomycin, 10 ng/mL of IL-3, and 30 ng/mL of Stem Cell Factor. The isolated cells were further cultured at 37 °C in 5% CO_2_. Two days after isolation, the culture medium was replaced with fresh medium. On day nine, the cells were detached from the flask bottom with gentle pipetting and further cultured at a concentration of 1×10^6^ cells/mL in fresh medium in a new flask. The PMCs were used for the experiments on days 12-16 after isolation. For the antigen-stimulation experiments, the cells were treated overnight with 300 ng/mL anti-Dinitrophenyl IgE antibodies.

For Ca^2+^ imaging experiments, PMCs were loaded in a salt solution containing 2.5 µM Fura-2 AM and 0.1% Pluronic F-127 for 30 min at RT on the coverslips covered with concanavalin A (0.1 mg/mL). The intracellular free Ca^2+^ concentration was measured on the stage of an AxioObserver-A3 inverted microscope (Zeiss, Germany) equipped with a 40x (1.3NA) immersion oil objective (Zeiss, Germany) in a perfused 0.5 mL chamber at RT. The Physiological Salt Solution (PSS) contained (in mM): NaCl 135; KCl 6, CaCl_2_ 2, MgCl_2_ 1.2, glucose 12, and HEPES 10; pH 7.4 (NaOH). At the beginning of each experiment, the cells were washed thoroughly with PSS. The fluorescence signal was obtained by alternately exciting Fura-2 with 340 nm and 380 nm light (50 ms exposure time) using a pE-800fura (CoolLED, United Kingdom) light source. The emitted fluorescent signal was filtered for wavelengths >510 nm and detected with an AxioCam 503 mono CCD camera (Zeiss, Germany). The fluorescent ratio F340/F380 was measured with an acquisition rate of 5 s per cycle. The camera and the Light source were controlled by the Zen 3.2 software (Zeiss, Germany).

### Isolation of primary T cells and microdomain Ca^2+^ measurements

Fura2-AM, Fluo4-AM, and Fura-Red-AM were obtained from Life Technologies, dissolved in dimethyl sulfoxide, and stored at −20°C. Anti-mouse CD3ε and anti-mouse CD28 mAb monoclonal antibodies were obtained from BD Biosciences. CD4^+^ T cells were isolated from spleens by negative selection using the EasySep Mouse CD4^+^ T cell Enrichment Kit (STEMCELL Technologies Inc.) according to the manufacturer’s instructions.

For global Ca^2+^ imaging, T cells were loaded with 4µM Fura2-AM for 30 min at 37°C, washed twice in imaging buffer [140 mM NaCl, 5 mM KCl, 1 mM MgSO4, 1 mM CaCl_2_, 20 mM Hepes (pH 7.4), 1 mM NaH_2_PO_4_, and 5 mM glucose], then kept at RT for 20 min for de-esterification. Cells were placed on BSA (5 mg/mL) / Poly-L-Lysine (0.1 mg/mL) coated slides and stimulated with 1 µg/mL anti-CD3ε for 60s. 1.67 µM thapsigargin was added at 720 s as a positive control (data not shown). Images were acquired on a Leica IRBE microscope with 40-fold magnification, a Sutter DG-4 light source, and an electron-multiplying charge-coupled device camera (Hamamatsu). One frame every 2 s was acquired using a Fura-2 filter set [excitation (ex), HC 340/26 nm, HC 387/11 nm; beam splitter (bs), 400DCLP; emission (em), 510/84 nm; AHF Analysentechnik]. Intracellular Ca^2+^ concentration was determined based on the equation of Grynkiewicz et al. 1985 (Grynkiewicz et al., 1985). Therefore, R_min_ [using the lowest ratio (R) and fluorescence (F) after EGTA chelation] and R_max_ (using the highest R and F after ionomycin incubation) of Fura-2 in single-cell measurements were assessed.

Imaging of Ca^2+^ microdomains in Fluo4- and FuraRed-loaded T cells was carried out as described in Gerlach et al. 2024 (Gerlach et al., 2024). Imaging was carried out with an exposure time of 25 ms (40 frames/s) in a 14-bit mode using a dual-view module (Optical Insights, PerkinElmer Inc.) to split the emission wavelengths [filters: ex, 480/40; bs, 495; emission 1 (em1) 1, 542/50; em2, 650/57]. The DARTS pipeline for postprocessing and analysis was used (Woelk et al., 2023).

### RNA isolation from murine cardiomyocytes and spleen for qPCR

Total RNA of ventricular cardiomyocytes was extracted using the miRNeasy Micro Kit (Qiagen) combined with the RNase Free DNase Set (Qiagen). RNA quantity, purity, and integrity were assessed by spectrophotometry (TECAN) using A260/280 and A260/230 ratios.

For spleen tissue, total RNA was extracted using TRIzol reagent (Invitrogen, ThermoFisher Scientific, Germany) and a Kinematica Polytron homogenizer. Briefly, after homogenization and a 10 min incubation at RT, the samples were centrifuged at 13,000 rpm for 10 min at 4 °C. The cleared supernatant was retained for phase separation with chloroform (0.2 ml per 1 ml TRIzol), tubes were shaken vigorously for 15 s, and incubated 2 min at RT. After centrifugation at 13,000 rpm for 15 min at 4 °C, the upper aqueous phase was collected. RNA was precipitated by adding 0.5 ml 100 % isopropanol per 1 ml TRIzol, incubated 10 min at RT, and centrifuged at 13,000 rpm for 10 min at 4 °C. Pellets were washed twice with 75 % ice-cold ethanol, with brief vortexing and centrifugation at ∼10,800 rpm for 5 min at 4 °C. Ethanol was removed. Pellets were air-dried for 10 min at RT. RNA was resuspended in 50 µl RNase-free water and incubated for 10 min at 55 °C. An additional DNase digestion step was performed with the TURBO DNA-free Kit (Invitrogen). RNA was quantified by spectrophotometry (TECAN) using A260/280 and A260/230 ratios before and after DNase treatment.

Isolated RNA from both cardiomyocytes and spleen was used for cDNA synthesis from 1 µg total RNA with the SensiFAST cDNA Synthesis kit (meridian Bioscience), followed by Real-time quantitative PCR using SYBR Master Mix.

### Statistical analysis

Results were analyzed using OriginPro 2019 (OriginLab) and Graphpad Prism 8.0. Values are shown as mean ± SEM. Statistical significances were assessed using the appropriate statistical tests (unpaired Student’s t-test, Mann-Whitney U test, or Two-Way ANOVA). Significance is depicted as * p < 0.05; ** p < 0.01, *** p < 0.001.

## Acknowledgements

This research was funded by the German Research Foundation (DFG) through the Collaborative Research Centre CRC1550 (project number FKZ 464424253), project B09 to MF; SFB1328 (project number FKZ 335447717), project A01 to AHG, and project A21 to MF; FOR 5705 – Neuroflame project P03 (Project number 523862973, FR 1638/6-1) to MF and AHG. MD student Jung Fellowship to KJW from the Jung Stiftung für Wissenschaft und Forschung (Hamburg, Germany), the DZHK (German Centre for Cardiovascular Research), Heidelberg/Mannheim, and the BMBF (German Ministry of Education and Research).

## Supplementary figures

**Suppl. Fig. 1.**
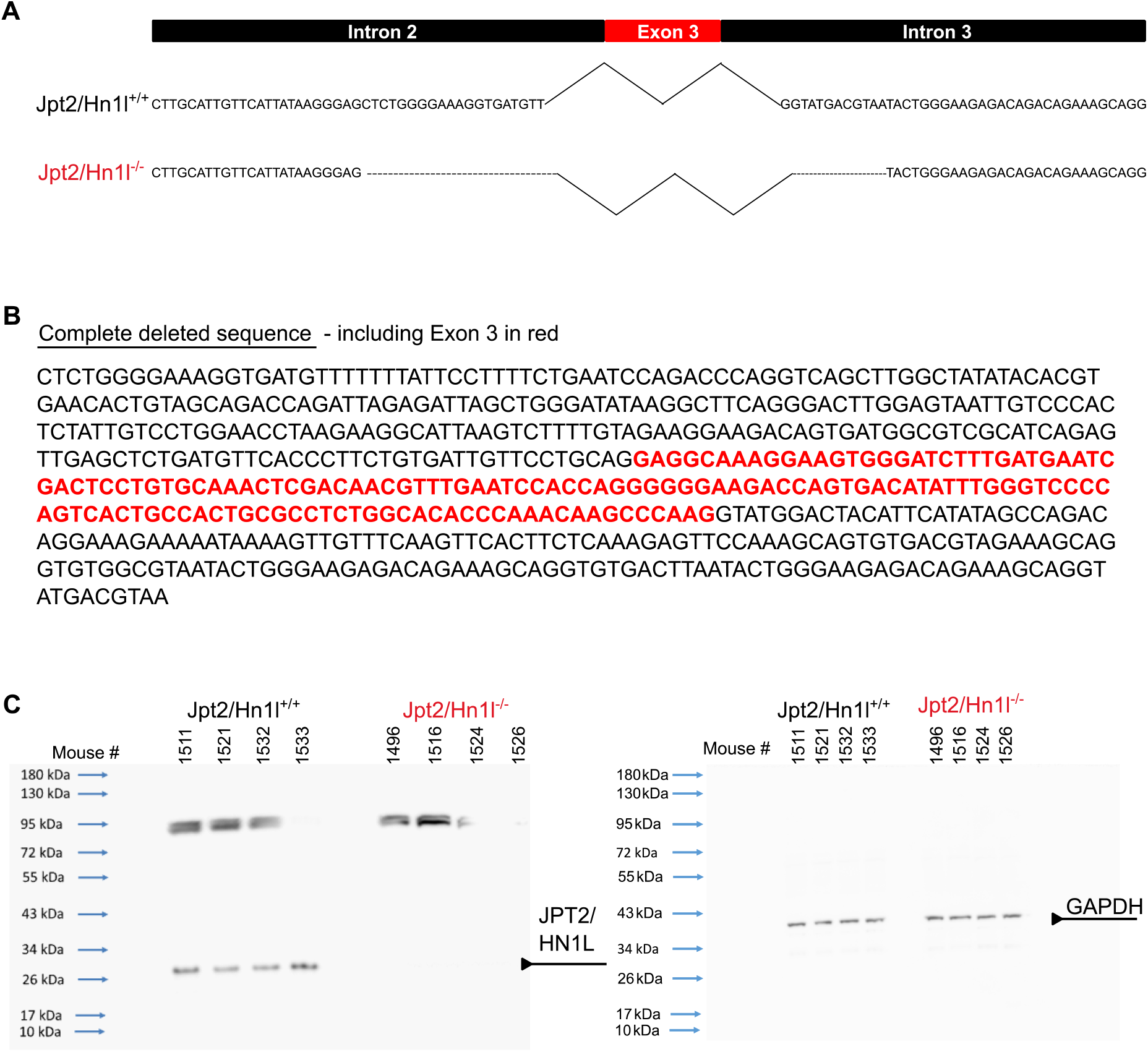
Genetic deletion of Jpt2/Hn1l in mice. **(A)** We generated a Jpt2/Hn1l null allele by deletion of exon 3. **(B)** The complete sequence that was removed from the genome was 577 bp long, containing the 143 bp long exon 3 (red). **(C)** We confirmed by Western Blot analysis that the 26/27 kDa Protein was not expressed in our Jpt2/Hn1l–/– mice and used GAPDH as a housekeeping protein.

**Suppl. Fig. 2.**
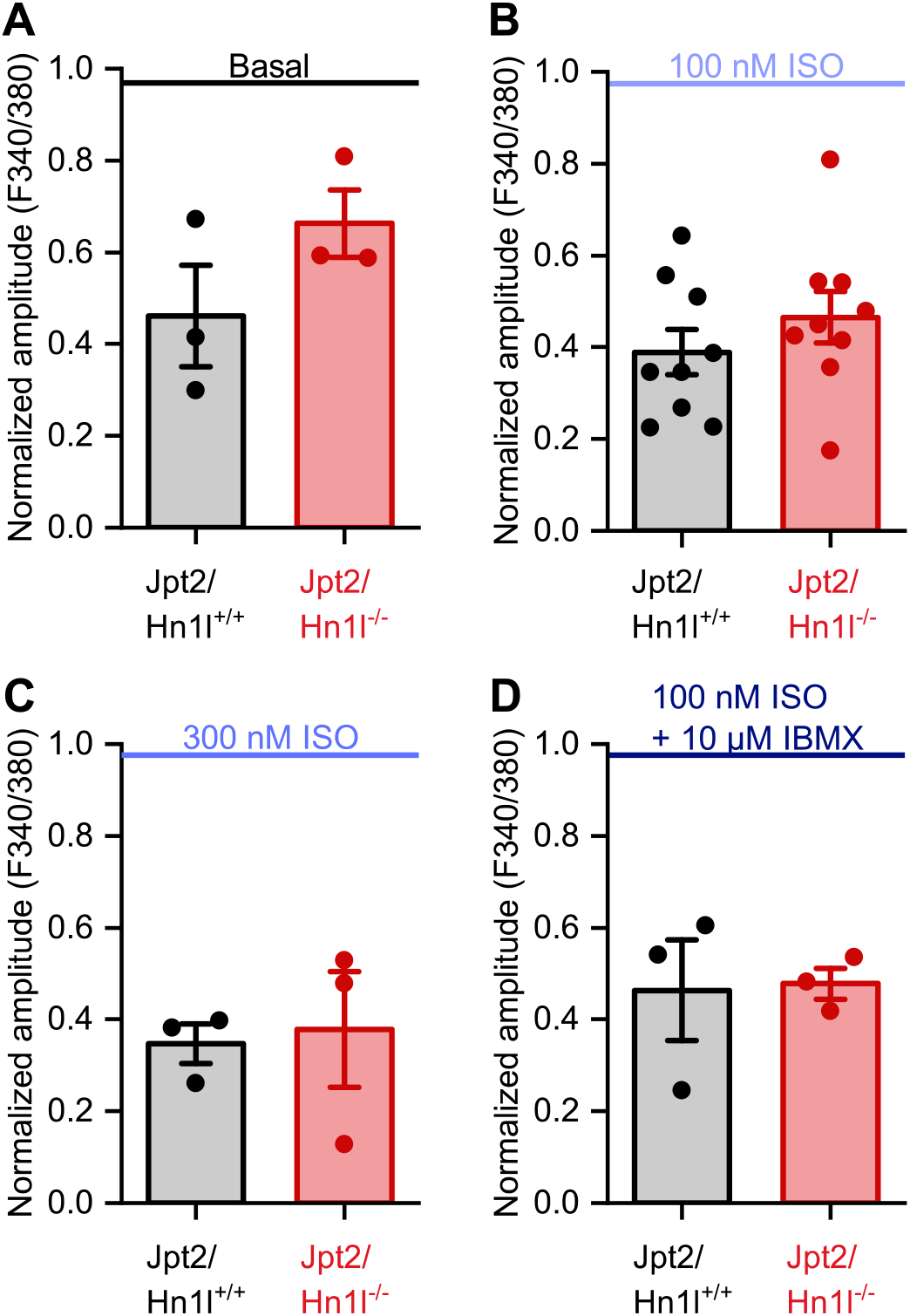
Amplitude analysis of paced cardiomyocytes. Analysis of amplitude of Ca^2+^ transients evoked by **(A)** pacing alone or in combination with **(B)** 100nM ISO, **(C)** 300nM ISO, or **(D)** a combination of 100nM ISO and 10µM IBMX.

**Suppl. Fig. 3.**
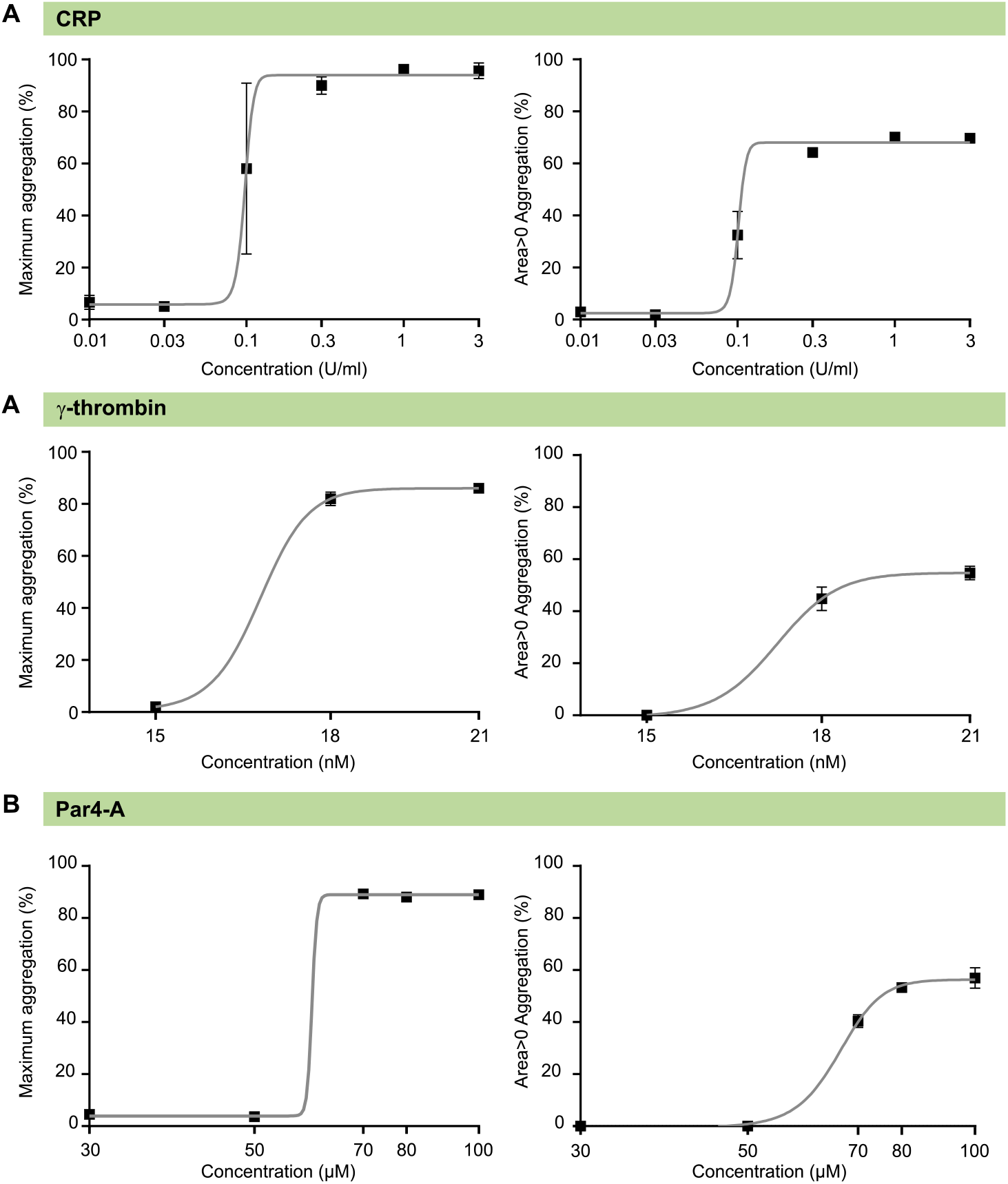
Dose-response curve of aggregation of WT platelets. **(A-B)** Dose response curves of aggregation evoked by **(A)** γ-thrombin, or **(B)** PAR4-A peptide.

**Suppl. Fig. 4.**
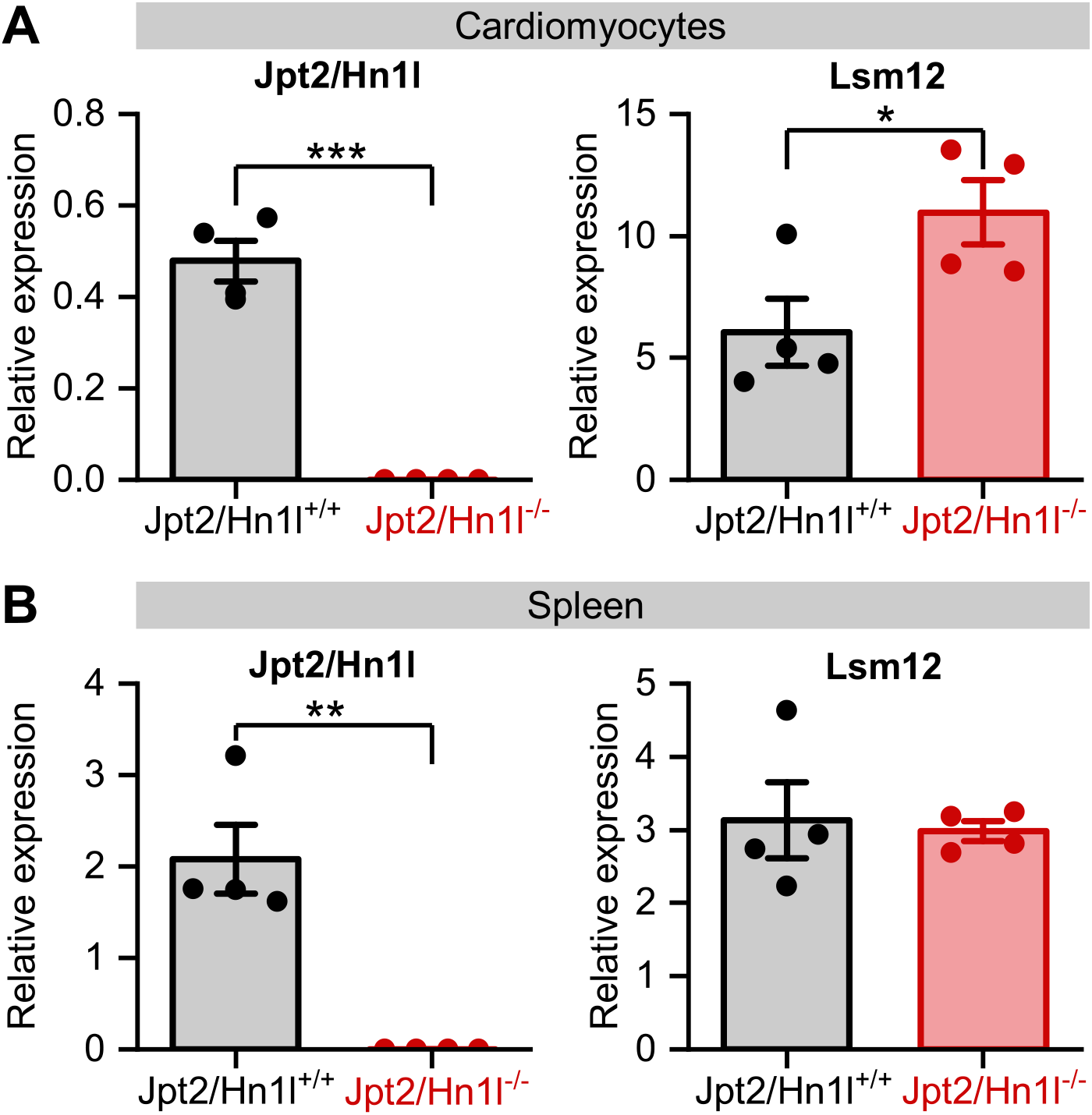
Relative mRNA expression by qPCR. Relative expression levels of Jpt2/Hn1l and Lsm12 in **(A)** cardiomyocytes and **(B)** spleens of Jpt2/Hn1l^+/+^ and Jpt2/Hn1l^−/−^ mice.

## References

Arlt, E., Fraticelli, M., Tsvilovskyy, V., Nadolni, W., Breit, A., O’Neill, T. J., Resenberger, S., Wennemuth, G., Wahl-Schott, C., Biel, M., Grimm, C., Freichel, M., Gudermann, T., Klugbauer, N., Boekhoff, I., & Zierler, S. (2020). TPC1 deficiency or blockade augments systemic anaphylaxis and mast cell activity. Proceedings of the National Academy of Sciences of the United States of America, 117(30), 18068–18078. 10.1073/pnas.1920122117

Berg, I., Potter, B. V, Mayr, G. W., & Guse, A. H. (2000). Nicotinic acid adenine dinucleotide phosphate (NAADP(+)) is an essential regulator of T-lymphocyte Ca(2+)-signaling. The Journal of Cell Biology, 150(3), 581–588. 10.1083/jcb.150.3.581

Calcraft, P. J., Arredouani, A., Ruas, M., Pan, Z., Cheng, X., Hao, X., Tang, J., Rietdorf, K., Teboul, L., Chuang, K., Lin, P., Xiao, R., Wang, C., Zhu, Y., Lin, Y., Wyatt, C. N., Parrington, J., Ma, J., Evans, A. M., … Zhu, M. X. (2009). NAADP mobilizes calcium from acidic organelles through two-pore channels. 459(7246), 596–600. 10.1038/nature08030.NAADP

Camacho Londoño, J. E., Tian, Q., Hammer, K., Schröder, L., Camacho Londoño, J., Reil, J. C., He, T., Oberhofer, M., Mannebach, S., Mathar, I., Philipp, S. E., Tabellion, W., Schweda, F., Dietrich, A., Kaestner, L., Laufs, U., Birnbaumer, L., Flockerzi, V., Freichel, M., & Lipp, P. (2015). A background Ca2+ entry pathway mediated by TRPC1/TRPC4 is critical for the development of pathological cardiac remodelling. European Heart Journal, 36(33), 2257–2266. 10.1093/eurheartj/ehv250

Capel, R. A., Bolton, E. L., Lin, W. K., Aston, D., Wang, Y., Liu, W., Wang, X., Burton, R. A. B., Bloor-Young, D., Shade, K. T., Ruas, M., Parrington, J., Churchill, G. C., Lei, M., Galione, A., & Terrar, D. A. (2015). Two-pore channels (TPC2s) and nicotinic acid adenine dinucleotide phosphate (NAADP) at lysosomal-sarcoplasmic reticular junctions contribute to acute and chronic β-adrenoceptor signaling in the heart. Journal of Biological Chemistry, 290(50), 30087–30098. 10.1074/jbc.M115.684076

Capel, R. A., & Terrar, D. A. (2015). The importance of Ca2+-dependent mechanisms for the initiation of the heartbeat. Frontiers in Physiology, 6(MAR), 1–19. 10.3389/fphys.2015.00080

Castonguay, J., Orth, J. H. C., Müller, T., Sleman, F., Grimm, C., Wahl-Schott, C., Biel, M., Mallmann, R. T., Bildl, W., Schulte, U., & Klugbauer, N. (2017). The two-pore channel TPC1 is required for efficient protein processing through early and recycling endosomes. Scientific Reports, 7(1), 1–15. 10.1038/s41598-017-10607-4

Chang, F. S., Wang, Y., Dmitriev, P., Gross, J., Galione, A., & Pears, C. (2020). A two-pore channel protein required for regulating mTORC1 activity on starvation. BMC Biology, 18(1), 1–16. 10.1186/s12915-019-0735-4

Chen, X., Song, Q. L., Ji, R., Wang, J. Y., Cao, M. L., Guo, D. Y., Zhang, Y., & Yang, J. (2024). JPT2 Affects Trophoblast Functions and Macrophage Polarization and Metabolism, and Acts as a Potential Therapeutic Target for Recurrent Spontaneous Abortion. Advanced Science, 11(16), 1–17. 10.1002/advs.202306359

Churamani, D., Carrey, E. A., Dickinson, G. D., & Patel, S. (2004). Determination of cellular nicotinic acid-adenine dinucleotide phosphate (NAADP) levels. Biochemical Journal, 380(2), 449–454. 10.1042/BJ20031754

Coxon, C. H., Lewis, A. M., Sadler, A. J., Vasudevan, S. R., Thomas, A., Dundas, K. A., Taylor, L., Campbell, R. D., Gibbins, J. M., Churchill, G. C., & Tucker, K. L. (2012). NAADP regulates human platelet function. Biochemical Journal, 441(1), 435–442. 10.1042/BJ20111175

Dammermann, W., Zhang, B., Nebel, M., Cordiglieri, C., Odoardi, F., Kirchberger, T., Kawakami, N., Dowden, J., Schmid, F., Dornmair, K., Hohenegger, M., Flügel, A., Guse, A. H., & Potter, B. V. L. (2009). NAADP-mediated Ca2+ signaling via type 1 ryanodine receptor in T cells revealed by a synthetic NAADP antagonist. Proceedings of the National Academy of Sciences of the United States of America, 106(26), 10678–10683. 10.1073/pnas.0809997106

Davidson, S. M., Foote, K., Kunuthur, S., Gosain, R., Tan, N., Tyser, R., Zhao, Y. J., Graeff, R., Ganesan, A., Duchen, M. R., Patel, S., & Yellon, D. M. (2015). Inhibition of NAADP signalling on reperfusion protects the heart by preventing lethal calcium oscillations via two-pore channel 1 and opening of the mitochondrial permeability transition pore. Cardiovascular Research, 108(3), 357–366. 10.1093/cvr/cvv226

Diercks, B.-P., Werner, R., Weidemüller, P., Czarniak, F., Hernandez, L., Lehmann, C., Rosche, A., Krüger, A., Kaufmann, U., Vaeth, M., Failla, A. V, Zobiak, B., Kandil, F. I., Schetelig, D., Ruthenbeck, A., Meier, C., Lodygin, D., Flügel, A., Ren, D., … Guse, A. H. (2018). ORAI1, STIM1/2, and RYR1 shape subsecond Ca2+ microdomains upon T cell activation. Science Signaling, 11(561), 1–14. 10.1126/scisignal.aat0358

Feng, M., Elaïb, Z., Borgel, D., Denis, C. V., Adam, F., Bryckaert, M., Rosa, J. P., & Bobe, R. (2020). NAADP/SERCA3-Dependent Ca2+Stores Pathway Specifically Controls Early Autocrine ADP Secretion Potentiating Platelet Activation. Circulation Research, 127(7), e166–e183. 10.1161/CIRCRESAHA.119.316090

Galione, A. (2011). NAADP Receptors. Cold Spring Harbor Perspectives in Biology, 3(1), 1–17. 10.1101/cshperspect.a004036

Gasser, A., Bruhn, S., & Guse, A. H. (2006). Second messenger function of nicotinic acid adenine dinucleotide phosphate revealed by an improved enzymatic cycling assay. Journal of Biological Chemistry, 281(25), 16906–16913. 10.1074/jbc.M601347200

Gerasimenko, J. V., Charlesworth, R. M., Sherwood, M. W., Ferdek, P. E., Mikoshiba, K., Parrington, J., Petersen, O. H., & Gerasimenko, O. V. (2015). Both RyRs and TPCs are required for NAADP-induced intracellular Ca2+ release. Cell Calcium, 58(3), 237–245. 10.1016/j.ceca.2015.05.005

Gerlach, F., Möckl, F., Kovacevic, D., Brock, V. J., Winzer, R., Meyer, L., Lohr, D., Woelk, L. M., Tolosa, E., Werner, R., & Diercks, B. P. (2024). Imaging Initial Ca2+ Microdomains in Primary T Cells. Journal of Visualized Experiments : JoVE, 212, 1–16. 10.3791/67075

Golde, W. T., Gollobin, P., & Rodriguez, L. L. (2005). A rapid, simple, and humane method for submandibular bleeding of mice using a lancet. Lab Animal, 34(9), 39–43. 10.1038/laban1005-39

Grynkiewicz, G., Poenie, M., & Tsien, R. Y. (1985). A new generation of Ca2+ indicators with greatly improved fluorescence properties. Journal of Biological Chemistry, 260(6), 3440–3450. 10.1016/s0021-9258(19)83641-4

Gu, F., Krüger, A., Roggenkamp, H. G., Alpers, R., Lodygin, D., Jaquet, V., Möckl, F., Hernandez C L. C., Winterberg, K., Bauche, A., Rosche, A., Grasberger, H., Kao, J. Y., Schetelig, D., Werner, R., Schröder, K., Carty, M., Bowie, A. G., Huber, S., … Guse, A. H. (2021). Dual NADPH oxidases DUOX1 and DUOX2 synthesize NAADP and are necessary for Ca2+ signaling during T cell activation. Science Signaling, 14(709), eabe3800. 10.1126/scisignal.abe3800

Gunaratne, G. S., Brailoiu, E., He, S., Unterwald, E. M., Patel, S., Slama, J. T., Walseth, T. F., & Marchant, J. S. (2021). Essential requirement for JPT2 in NAADP-evoked Ca2+signaling. Science Signaling, 14(675). 10.1126/scisignal.abd5605

Gunaratne, G. S., Brailoiu, E., Kumar, S., Yuan, Y., Slama, J. T., Walseth, T. F., Patel, S., & Marchant, J. S. (2023). Convergent activation of two-pore channels mediated by the NAADP-binding proteins JPT2 and LSM12. Science Signaling, 16(799), 1–24. 10.1126/scisignal.adg0485

Guse, A. H., & Wolf, I. M. A. (2016). Ca2+ microdomains, NAADP, and type 1 ryanodine receptor in cell activation. Biochimica et Biophysica Acta - Molecular Cell Research, 1863(6), 1379–1384. 10.1016/j.bbamcr.2016.01.014

Hermann, J., Bender, M., Schumacher, D., Woo, M. S., Shaposhnykov, A., Rosenkranz, S. C., Kuryshev, V., Meier, C., Guse, A. H., Friese, M. A., Freichel, M., & Tsvilovskyy, V. (2020). Contribution of NAADP to Glutamate-Evoked Changes in Ca2+ Homeostasis in Mouse Hippocampal Neurons. Frontiers in Cell and Developmental Biology, 8(June), 1–13. 10.3389/fcell.2020.00496

Heßling, L. D., Troost-Kind, B., & Weiß, M. (2023). NAADP-binding proteins — Linking NAADP signaling to cancer and immunity. Biochimica et Biophysica Acta - Molecular Cell Research, 1870(7). 10.1016/j.bbamcr.2023.119531

Hohenegger, M., Suko, J., Gscheidlinger, R., Drobny, H., & Zidar, A. (2002). Nicotinic acid-adenine dinucleotide phosphate activates the skeletal muscle ryanodine receptor. The Biochemical Journal, 367(Pt 2), 423–431. 10.1042/BJ20020584

Jardin, I., Ben Amor, N., Bartegi, A., Pariente, J. A., Salido, G. M., & Rosado, J. A. (2007). Differential involvement of thrombin receptors in Ca2+ release from two different intracellular stores in human platelets. Biochemical Journal, 401(1), 167–174. 10.1042/BJ20060888

Krukenberg, S., Möckl, F., Weiß, M., Dekiert, P., Hofmann, M., Gerlach, F., Winterberg, K. J., Kovacevic, D., Khansahib, I., Troost, B., Hinrichs, M., Granato, V., Nawrocki, M., Hub, T., Tsvilovskyy, V., Medert, R., Woelk, L. M., Förster, F., Li, H., … Guse, A. H. (2024). MASTER-NAADP: a membrane permeable precursor of the Ca2+ mobilizing second messenger NAADP. Nature Communications, 15(1). 10.1038/s41467-024-52024-y

Lin-Moshier, Y., Keebler, M. V., Hooper, R., Boulware, M. J., Liu, X., Churamani, D., Abood, M. E., Walseth, T. F., Brailoiu, E., Patel, S., & Marchant, J. S. (2014). The Two-pore channel (TPC) interactome unmasks isoform-specific roles for TPCs in endolysosomal morphology and cell pigmentation. Proceedings of the National Academy of Sciences of the United States of America, 111(36), 13087–13092. 10.1073/pnas.1407004111

Lin, W. K., Bolton, E. L., Cortopassi, W. A., Wang, Y., O’Brien, F., Maciejewska, M., Jacobson, M. P., Garnham, C., Ruas, M., Parrington, J., Lei, M., Sitsapesan, R., Galione, A., & Terrar, D. A. (2017). Synthesis of the Ca2+-mobilizing messengers NAADP and cADPR by intracellular CD38 enzyme in the mouse heart: Role in -adrenoceptor signaling. Journal of Biological Chemistry, 292(32), 13243–13257. 10.1074/jbc.M117.789347

Medert, R., Pironet, A., Bacmeister, L., Segin, S., Londoño, J. E. C., Vennekens, R., & Freichel, M. (2020). Genetic background influences expression and function of the cation channel TRPM4 in the mouse heart. Basic Research in Cardiology, 115(6), 1–16. 10.1007/s00395-020-00831-x

Medert, R., Thumberger, T., Tavhelidse-Suck, T., Hub, T., Kellner, T., Oguchi, Y., Dlugosz, S., Zimmermann, F., Wittbrodt, J., & Freichel, M. (2023). Efficient single copy integration via homology-directed repair (scHDR) by 5′modification of large DNA donor fragments in mice. Nucleic Acids Research, 51(3), E14. 10.1093/nar/gkac1150

Morgan, A. J., & Galione, A. (2014). Two-pore channels (TPCs): Current controversies. BioEssays, 36(2), 173–183. 10.1002/bies.201300118

Morgan, A. J., Platt, F. M., Lloyd-Evans, E., & Galione, A. (2011). Molecular mechanisms of endolysosomal Ca 2+ signalling in health and disease. Biochemical Journal, 439(3), 349–378. 10.1042/BJ20110949

Naylor, E., Arredouani, A., Vasudevan, S. R., Lewis, A. M., Parkesh, R., Mizote, A., Rosen, D., Thomas, J. M., Izumi, M., Ganesan, A., Galione, A., & Churchill, G. C. (2009). Identification of a chemical probe for NAADP by virtual screening. Nature Chemical Biology, 5(4), 220–226. 10.1038/nchembio.150

Nebel, M., Schwoerer, A. P., Warszta, D., Siebrands, C. C., Limbrock, A. C., Swarbrick, J. M., Fliegert, R., Weber, K., Bruhn, S., Hohenegger, M., Geisler, A., Herich, L., Schlegel, S., Carrier, L., Eschenhagen, T., Potter, B. V. L., Ehmke, H., & Guse, A. H. (2013). Nicotinic acid adenine dinucleotide phosphate (NAADP)-mediated calcium signaling and arrhythmias in the heart evoked by β-adrenergic stimulation. Journal of Biological Chemistry, 288(22), 16017–16030. 10.1074/jbc.M112.441246

Roggenkamp, H. G., Khansahib, I., Hernandez C L. C., Zhang, Y., Lodygin, D., Krüger, A., Gu, F., Möckl, F., Löhndorf, A., Wolters, V., Woike, D., Rosche, A., Bauche, A., Schetelig, D., Werner, R., Schlüter, H., Failla, A. V, Meier, C., Fliegert, R., … Guse, A. H. (2021). HN1L/JPT2: A signaling protein that connects NAADP generation to Ca2+ microdomain formation. Science Signaling, 14(675). 10.1126/scisignal.abd5647

Rosado, J. A. (2011). Acidic Ca2+ stores in platelets. Cell Calcium, 50(2), 168–174. 10.1016/j.ceca.2010.11.011

Ruas, M., Davis, L. C., Chen, C.-C., Morgan, A. J., Chuang, K.-T., Walseth, T. F., Grimm, C., Garnham, C., Powell, T., Platt, N., Platt, F. M., Biel, M., Wahl-Schott, C., Parrington, J., & Galione, A. (2015). Expression of Ca2+-permeable two-pore channels rescues NAADP signalling in TPC-deficient cells. The EMBO Journal, 34(13), 1743–1758. 10.15252/embj.201490009

Soslau, G., Goldenberg, S. J., Class, R., & Jameson, B. (2004). Differential activation and inhibition of human platelet thrombin receptors by structurally distinct α-, β- and γ-thrombin. Platelets, 15(3), 155–166. 10.1080/0953710042000199848

Tabrizi, S. D., Nawrocki, M., Diercks, B.-P., Bedke, T., Böttcher, M., Stumme, F., Birus, M., Möckl, F., Hernandez, L., Gagliani, N., Lodygin, D., Flügel, A., Guse, A., Mittrücker, H.-W., & Huber, S. (2026). Ryanodine receptor 1 is dispensable for CD4 + T-cell differentiation and effector function in intestinal inflammation models. 10.64898/2026.01.14.699480

Thumberger, T., Tavhelidse-Suck, T., Gutierrez-Triana, J. A., Cornean, A., Medert, R., Welz, B., Freichel, M., & Wittbrodt, J. (2022). Boosting targeted genome editing using the hei-tag. ELife, 11, 1–16. 10.7554/eLife.70558

Woelk, L. M., Kovacevic, D., Husseini, H., Förster, F., Gerlach, F., Möckl, F., Altfeld, M., Guse, A. H., Diercks, B. P., & Werner, R. (2023). DARTS: an open-source Python pipeline for Ca2+ microdomain analysis in live cell imaging data. Frontiers in Immunology, 14(January), 1–11. 10.3389/fimmu.2023.1299435

Wolf, I. M. A., Diercks, B. P., Gattkowski, E., Czarniak, F., Kempski, J., Werner, R., Schetelig, D., Mittrücker, H. W., Schumacher, V., Von Osten, M., Lodygin, D., Flügel, A., Fliegert, R., & Guse, A. H. (2015). Frontrunners of T cell activation: Initial, localized Ca2+ signals mediated by NAADP and the type 1 ryanodine receptor. Science Signaling, 8(398), 1–13. 10.1126/scisignal.aab0863

Yang, L., Ottenheijm, R., Worley, P., Freichel, M., & Camacho Londoño, J. E. (2022). Reduction in SOCE and Associated Aggregation in Platelets from Mice with Platelet-Specific Deletion of Orai1. Cells, 11(20), 1–18. 10.3390/cells11203225

Zhang, F., Jin, S., Yi, F., & Li, P. L. (2009). TRP-ML1 functions as a lysosomal NAADP-sensitive Ca 2+ release channel in coronary arterial myocytes. Journal of Cellular and Molecular Medicine, 13(9 B), 3174–3185. 10.1111/j.1582-4934.2008.00486.x

Zhang, F., Xu, M., Han, W. Q., & Li, P. L. (2011). Reconstitution of lysosomal NAADP-TRP-ML1 signaling pathway and its function in TRP-ML1 -/-cells. American Journal of Physiology - Cell Physiology, 301(2), 421–430. 10.1152/ajpcell.00393.2010

Zhang, J., Guan, X., Shah, K., & Yan, J. (2021). Lsm12 is an NAADP receptor and a two-pore channel regulatory protein required for calcium mobilization from acidic organelles. Nature Communications, 12(1), 1–13. 10.1038/s41467-021-24735-z

Zong, X., Schieder, M., Cuny, H., Fenske, S., Gruner, C., Rötzer, K., Griesbeck, O., Harz, H., Biel, M., & Wahl-Schott, C. (2009). The two-pore channel TPCN2 mediates NAADP-dependent Ca2+-release from lysosomal stores. Pflugers Archiv European Journal of Physiology, 458(5), 891–899. 10.1007/s00424-009-0690-y

